# Prot2Prot: A Deep Learning Model for Rapid, Photorealistic Macromolecular Visualization

**DOI:** 10.1101/2022.03.21.485218

**Authors:** Jacob D. Durrant

## Abstract

**Motivation:** Molecular visualization is a cornerstone of structural biology, providing insights into the form and function of biomolecules that are difficult to achieve any other way. Scientific analysis, publication, education, and outreach often benefit from photorealistic molecular depictions rendered using advanced computer-graphics programs such as Maya, 3ds Max, and Blender. However, setting up molecular scenes in these programs can be laborious even for expert users, and rendering often requires substantial time and computer resources.

**Results:** We have created a deep-learning model called Prot2Prot that quickly imitates photorealistic visualization styles, given a much simpler, easy-to-generate molecular representation. The resulting images are often indistinguishable from images rendered using industry-standard 3D graphics programs, but they can be created in a fraction of the time, even when running in a web browser. To the best of our knowledge, Prot2Prot is the first example of image-to-image translation applied to macromolecular visualization.

**Availability:** Prot2Prot is available free of charge, released under the terms of the Apache License, Version 2.0. Users can access a Prot2Prot-powered web app without registration at http://durrantlab.com/prot2prot.

## Introduction

Molecular visualization is a critical structural-biology tool that provides valuable insights into the form and function of biomolecules. Artistically rendered molecular images can also inspire students and the public via education and outreach. Though well-known programs such as VMD [2], UCSF Chimera [3], and PyMOL [4] were conceived principally as analysis tools, these programs can also produce striking images. But they understandably lack many of the advanced rendering techniques commonly used in the video game and film industries.

Several computer-graphics programs incorporate industry-standard techniques, including Maya, 3ds Max, and Blender. Among these, Blender is notable because it is free and open source. However, none of these programs are designed for molecular visualization specifically. Several programs and plugins seek to address this shortcoming, including our BlendMol plugin [5], which allows users to easily import molecular models prepared via VMD into the Blender environment. Though BlendMol greatly simplifies photorealistic molecular rendering, it still requires a good understanding of Blender, a program with a notoriously steep learning curve. Rendering high-quality images and videos is also computationally intensive, further limiting use. We here describe a deep-learning model called Prot2Prot that imitates a Blender-rendered molecular image given a much simpler and easier-to-generate representation (“sketch”) of a protein surface. Prot2Prot outputs an image that is often indistinguishable from a BlendMol-based visualization in a fraction of the time, allowing image “rendering” even in a web browser. The success of the Prot2Prot approach demonstrates how machine learning can serve as a valuable tool for enhancing scientific communication, with potential applications to fields beyond molecular visualization. We release Prot2Prot free of charge under the terms of the Apache License, Version 2.0. Users can access a Prot2Prot-powered web app without registration at http://durrantlab.com/prot2prot.

## Materials and Methods

### Simplified Protein-Surface “Sketches”

We first developed a simple 2D molecular representation that is easy to generate, even in slow and memory-limited environments such as web browsers (Figure 1A). We represent each atom as a simple circle, sized according to the van der Waals radius and distance from the virtual “camera” (i.e., depth). Each circle is outlined in black to gray depending on its depth to emphasize atomic boundaries.

**Figure 1.**
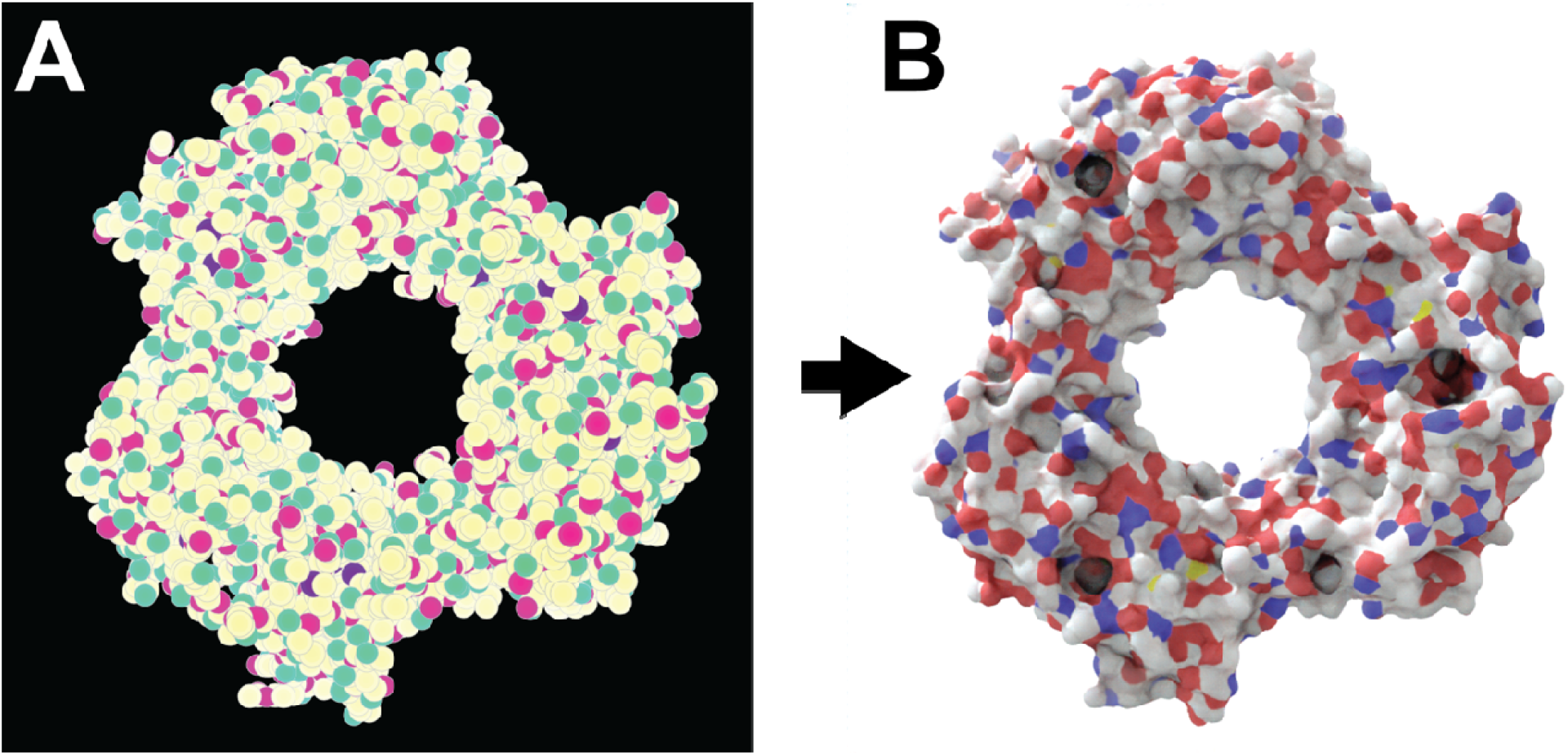
Prot2Prot image mapping. Proliferating cell nuclear antigen (PCNA) bound to the PCNA-interacting motif (PIP box) of the DNA-dependent metalloprotease SPRTN (DVC1; 6,099 atoms; PDB ID: 5IY4 [1]). A) The input image is a simplified 2D molecular representation that is trivial to generate. B) The output image mimics the appearance of a photorealistic rendering of the same protein, as if created using Blender/BlendMol.

The circles are colored according to carefully selected red, green, and blue (RGB) values. The red and green channels encode the atomic element. R/G values corresponding to carbon, nitrogen, oxygen, hydrogen, and phosphorus atoms are set at 100%/100%, 100%/0%, 0%/100%, 0%/50%, and 50%/50%, respectively. All other atoms are encoded as carbons. Depth is encoded on the blue (B) channel. It is set at 100% for those atoms close to the camera and 0% for those atoms that are distant.

To provide data sufficient for training a machine-learning model, we generated hundreds of representative protein-surface sketches that capture multiple proteins at many different angles and distances.

### Blender/BlendMol-Rendered Molecular Visualizations

For each protein-surface sketch, we rendered a matching photorealistic image using Blender 3.0.0, an open-source computer-graphics toolset. We created custom Python scripts that load a PDB file into Blender using the BlendMol plugin [5]; automatically adjust the focal point of the Blender camera; create a fog-to-white effect; set the surface materials, lighting, and other parameters; and render a photorealistic image to disk using the Cycles path-tracing render engine.

### Training a Prot2Prot Model to Map Input Sketches to Rendered Images

We trained a deep-learning model, Pix2Pix [6], to translate molecular sketches into the corresponding photorealistic protein images (Figure 1; PyTorch Pix2Pix implementation available at GitHub [7]). In the context of this project, we call the model Prot2Prot rather than Pix2Pix. We used the default values for training, except we selected U-Net 128 as the generator architecture and used instance rather than batch normalization. For each of three photorealistic styles, we trained separate Prot2Prot models to generate 256×256, 512×512, and 1024×1024 output images, respectively. In all cases, we trained on roughly 1,000 sketch/render pairs for 1,000 epochs using the default initial learning rate and then for another 1,000 epochs as the learning rate decayed linearly to zero.

To augment the data set available for training, we scaled the images by ∼112% and then randomly cropped them at the original size (e.g., 256×256 images were scaled to 286×286 and then randomly cropped to produce 256×256 images). To allow the models to learn to mimic the consistent directional lighting of the rendered target images, we did not rotate or flip images, an otherwise common technique used for further data augmentation.

### Model Conversion for Use with Tensorflow.js

We exported the trained PyTorch models to the ONNX format using the *torch*.*onnx*.*export* function. We then converted the ONNX files to the TensorFlow SavedModel format using the TensorFlow Backend for ONNX [8]. Finally, we converted the SavedModel files to the TensorFlow.js graph-model format using the *tensorflowjs_converter* command [9]. In this last step, we also applied 1-byte affine quantization to multiple nodes, which substantially reduced the file size without substantial impact on image quality.

### Colorization

Prot2Prot produces images with consistent, predetermined color schemes. However, users can modify the color palette after inference, allowing some degree of customizability. A color-intensity matrix, *w*, determines how much a user-specified color influences various portions of the output image. The entries of the matrix range from 0.0 (no influence on the output image) to 1.0 (full influence).

The color-intensity matrix is calculated via element-wise multiplication of four image-derived matrices. First, to leave non-protein-surface regions unmodified, we convert the input “sketch” to a binary mask, where entries corresponding to the protein surface are set to 1.0, and other entries are set to 0.0 (Figure 2A). Second, to influence well-lit protein-surface regions more than shadowed areas, we separately convert the output “rendered” image to a grayscale matrix whose values correspond to averaged red, green, and blue values, scaled from 0.0 to 1.0 (Figure 2B). Third, to preserve the fade-to-white fog effect, we create a third matrix from the blue channel of the input sketch image, which encodes depth (i.e., distance from the virtual camera). The values of this matrix range from 0.0 (most distant) to 1.0 (closest; Figure 2C). Fourth, to allow the user to control the colorization effect’s global strength, we create a matrix with identical entries equal to a user-defined color-strength parameter (Figure 2D). After the element-wise multiplication of these four matrixes, the final matrix (Figure 2E) can be optionally blurred to remove any sharp edges, per the user-defined color-blend parameter.

**Figure 2.**
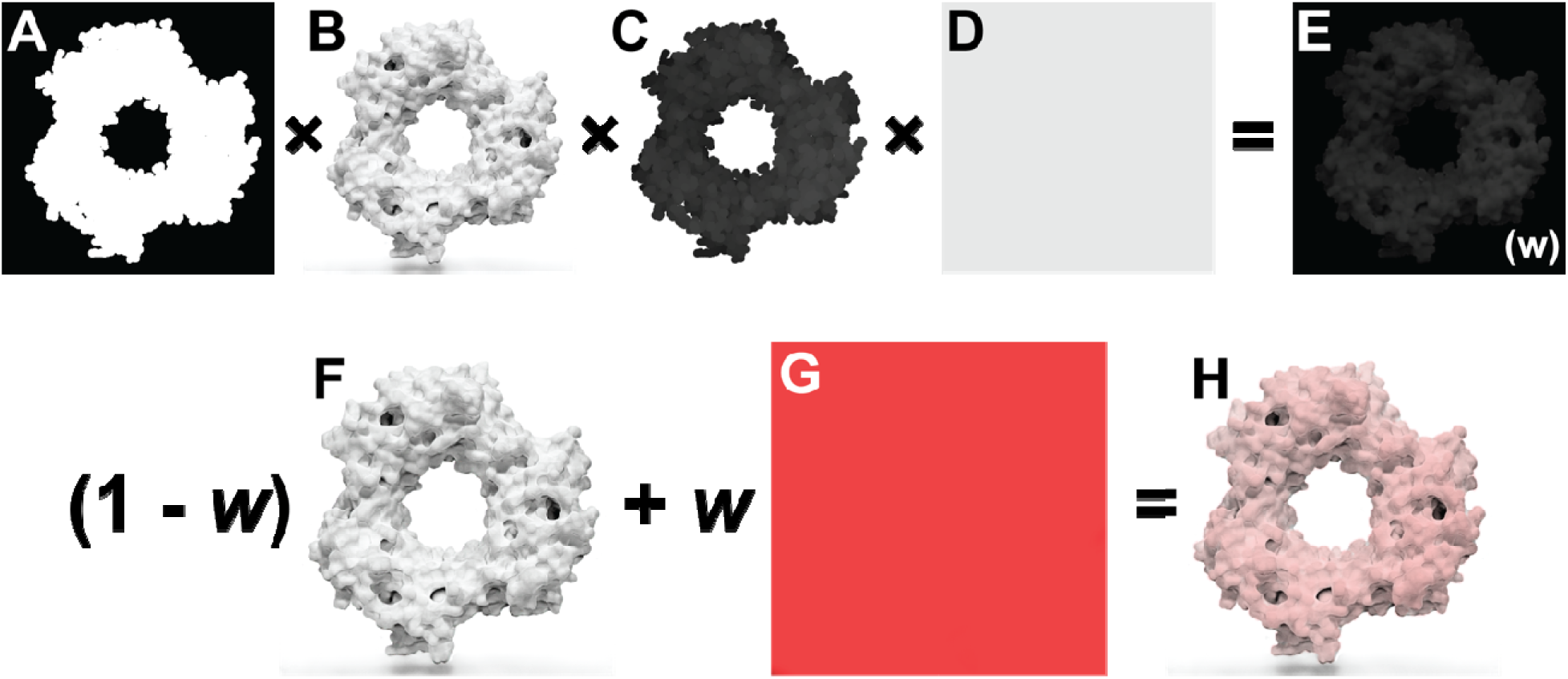
Prot2Prot colorization procedure. A) The mask matrix indicates which image regions include the protein surface. B) The grayscale matrix distinguishes well-lit and in-shadow protein-surface regions. C) The depth matrix indicates how far a protein region is from the virtual camera. D) The color-strength matrix allows the user to further modify the strength of the colorization effect. E) The final color-intensity matrix, called *w*, is calculated via element-wise multiplication of the four preceding matrices. F) The original Prot2Prot output image. G) A solid, user-specified color. H) The final image, created by averaging the images in F) and G), weighting by the color-intensity matrix, *w*.

We use this color-intensity matrix to adjust the original Prot2Prot output image (Figure 2F). A weighted average combines each pixel’s red, green, and blue values with those of a solid, user-defined color (Figure 2G). The pixel’s color is unchanged if the corresponding color-intensity-matrix value is 0.0 and replaced by the user-defined color entirely if the corresponding value is 1.0 (Figure 2H).

### Browser Implementation

We created a browser-based version of Prot2Prot following our established open-source approach [10–12]. The graphical user interface (GUI) is written in TypeScript using the Vue.js framework [13], the BootstrapVue CSS library [14], the TensorFlow.js machine-learning library [9], the Webpack module bundler [15], and Google’s Closure Compiler [16].

The “Input PDB File” panel (Figure 3A) allows users to load a PDB file into their web browser’s memory by selecting a file on their local computer or providing a PDB ID for remote download. Alternatively, clicking the “Use Example File” button automatically loads an example (PDB ID: 5IY4 [1]). Once the PDB file is loaded, the Prot2Prot user interface provides limited structure-editing options (e.g., users can remove ligands, water molecules, chains, etc.).

**Figure 3.**
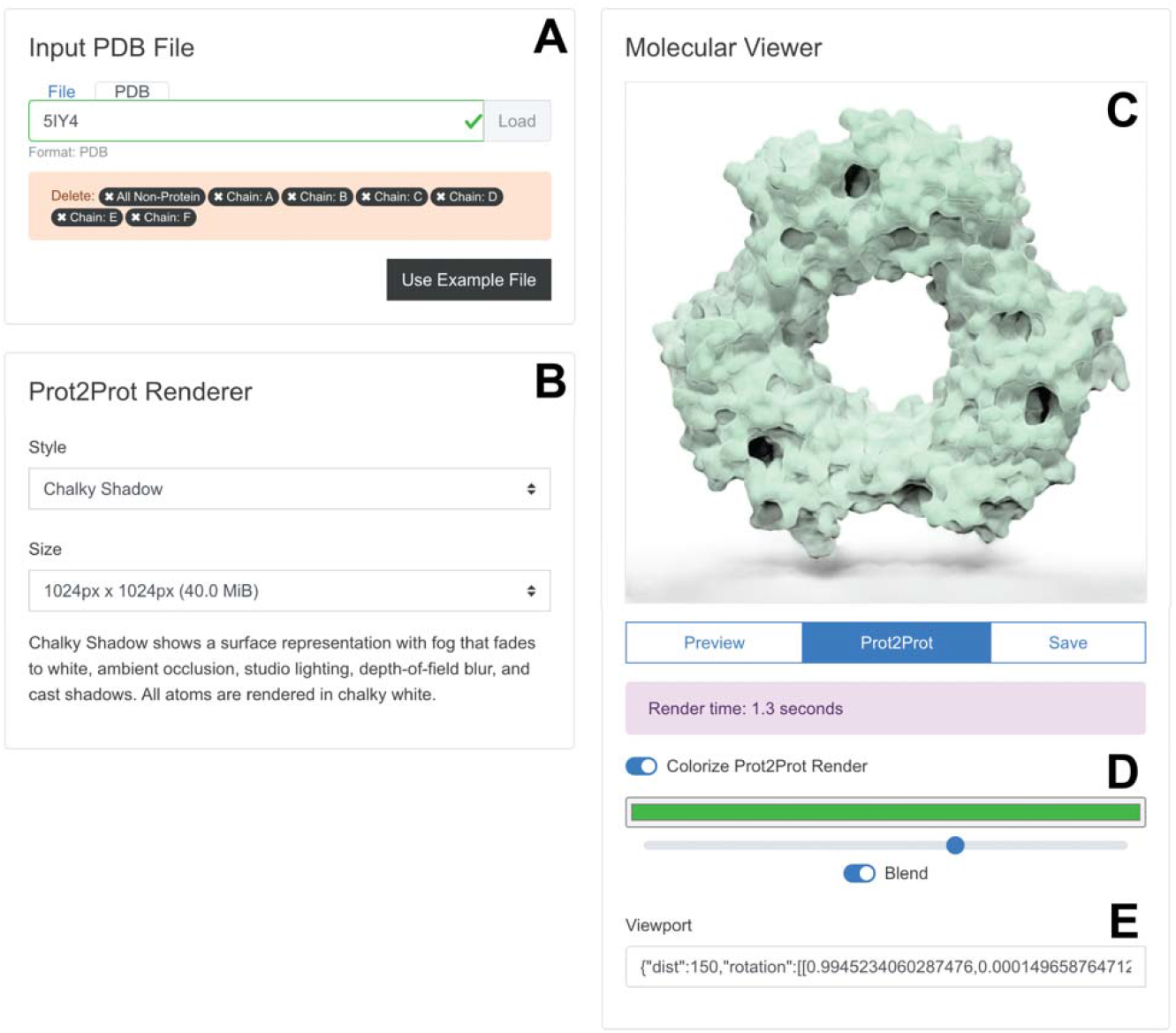
Browser-app user interface. A) The “Input PDB File” panel allows users to load and edit molecular structures. B) The “Prot2Prot Renderer” panel allows users to specify the rendering style and image size. C) The “Molecular Viewer” panel shows the rendered structure. D) Colorize options allow the user to adjust the protein color. E) The Viewport information can be copied and pasted to restore the rotation/zoom settings.

The “Prot2Prot Renderer” panel (Figure 3B) allows users to choose from various rendering styles and image dimensions (see Results and Discussion for a description). It also briefly explains the visual features of the selected style.

Users can position and display their molecules in the “Molecular Viewer” panel (Figure 3C). Structures are initially shown in “Preview” mode as fields of atomic spheres that can be easily rotated and scaled using the mouse, mouse wheel, or touch gestures. Once ready, the user clicks the “Prot2Prot” button to generate the corresponding photorealistic image in the browser. The “Save” button allows users to save the viewer image. Users can also toggle the “Colorize Prot2Prot Render” setting to specify color, color-strength, and color-blending options (Figure 3C, where green is selected). Finally, the app provides “Viewport” information that can be copied and pasted to restore the rotation/zoom settings (Figure 3D).

### Command-line-interface Implementation

Aside from running Prot2Prot in a web browser, users can also access the model via a command-line interface (CLI) powered by the Node.js JavaScript runtime environment. CLI Prot2Prot is well-suited for rendering single images and image sequences, which can be combined into videos. CLI Prot2Prot provides several default animations, including “still,” “rock,” “turntable” (rotation about a user-specified axis), and “zoom.” If a PDB file contains multiple frames, CLI Prot2Prot will also render protein dynamics, allowing users to visualize molecular dynamics simulations or interpolated protein structures (Video S1).

## Results and Discussion

The Prot2Prot machine-learning model effectively renders photorealistic molecular representations via image-to-image translation of a much simpler, easy-to-generate, molecular-surface “sketch.” Prot2Prot illustrations are well suited for scientific analysis, publication, outreach, and education. CLI Prot2Prot can also generate animations of protein motions (Videos S1 and S2).

### Description of Rendering Styles

We trained Prot2Prot models to mimic three distinctive rendering styles, which we call “Simple Surface,” “Chalky,” and “Chalky Shadow.”

### Simple Surface

In the “Simple Surface” rendering style, carbon, oxygen, nitrogen, sulfur, and hydrogen atoms are silver, red, blue, yellow, and white. Color support for other elements is limited. When rendering the photorealistic Blender images used for training, we applied two effects to give the final images a better sense of depth. First, we used Blender’s mist pass to render more distant protein regions in lighter colors, producing a “fade-to-white” fog effect. Second, we used Blender’s depth-of-field effect to focus the virtual camera on the protein surface directly in front of it, such that regions distant from that focal point appear blurred or out of focus.

We also used several advanced lighting techniques to enhance photorealism. First, we applied a slight subsurface-scattering effect to all surfaces using Blender’s Principled BSDF shader. When light hits many natural materials, it penetrates the surface and is scattered in the object’s interior. After a light ray makes its way back to the surface, it leaves the object at a random angle, not the predictable angle typical of a perfectly reflective (“glossy”) surface. Second, rather than light the scene with a single point or directional light, we used a public-domain, high dynamic range image (HDRI [17]) to surround and light the surfaces. High-dynamic-range (HDR) lighting prevents the darkest and lightest regions of the image from being saturated as perfectly black or white, allowing the viewer to see full detail across the entire image. Third, we also applied ambient occlusion to the scene. This non-physical rendering technique approximates global illumination by darkening surfaces that are only partially accessible to the broader environment (e.g., enclosed pockets). After rendering the image using Blender’s Cycles path-tracing render engine, we adjusted the color level using ImageMagick to ensure the background was precisely white, as typically required for publication-quality images. We successfully trained our Prot2Prot models to mimic these Blender-rendered output images given a corresponding input “sketch image.” When converted to the TensorFlow.js graph-model format, the final model takes up roughly 40 MB. Figure 4A and Figure 4B show how the model has learned to mimic the fade-to-white-fog (*), depth-of-field (†), and ambient-occlusion (‡) effects of the Blender-rendered training images.

**Figure 4.**
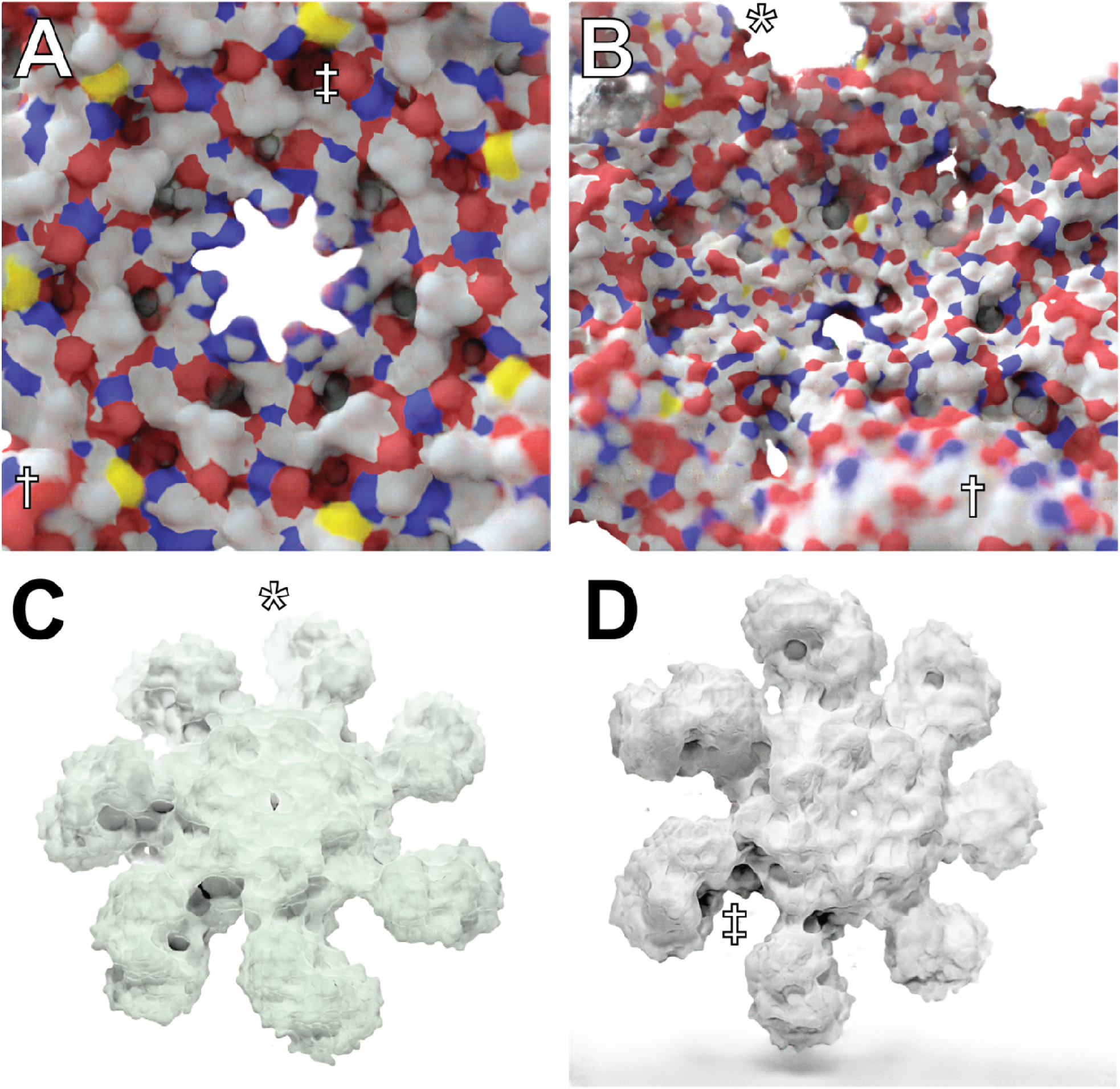
An atomic resolution model of the human apoptosome obtained via electron microscopy (70,189 atoms; PDB ID 3J2T), visualized using Prot2Prot. A) and B) Simple Surface rendering style. C) Chalky rendering style, colorized with a green tint. D) Chalky Shadow rendering style. Examples of fade-to-white fog, depth of field, and ambient occlusion are marked with *, †, and ‡, respectively.

### Chalky

The “Chalky” rendering style also has fade-to-white fog, ambient occlusion, and depth-of-field blur. Unlike Simple Surface, Chalky shows all atoms in the same white material, without subsurface scattering. Instead, we set the “Roughness” and “Clearcoat Roughness” settings on the Principled BSDF shader to their maximum values to give the surface a highly diffuse appearance. Chalky uses a public-domain studio lighting setup obtained from blendswap.com [18] to light the proteins rather than an HDRI. After rendering the training images, we again adjusted the color levels using ImageMagick.

Trained Prot2Prot models successfully mimic these Blender-rendered output images as well. The Chalky models also take up ∼40 MB, with similar run times in the browser. Figure 4C shows how Chalky images are particularly well suited to the custom colorization procedure (in this case, with a green tint) described in the Materials and Methods.

### Chalky Shadow

The “Chalky Shadow” rendering style is the same as the “Chalky” style, except the virtual studio lights are allowed to cast a shadow onto a pure-white floor below. The trained Prot2Prot models successfully mimic the shadows computed using advanced path tracing in Blender (Figure 4D). Video S2 (bottom row) illustrates how these shadows even convincingly change according to the protein orientation. These models are also roughly 40MB.

### Video Rendering via the Command Line Interface

Command-line-interface (CLI) Prot2Prot also accepts multi-frame PDB files as input, allowing users to create animations of molecular dynamics simulations, conformational transitions, etc. Prot2Prot provides four different animation styles via its CLI (Video S1). A “still” animation captures only the frame-by-frame motions of individual atoms without imparting any large-scale rotations to the entire protein. Alternatively, three whole-scene rotation animations can further facilitate visualization: “rock,” “turn table,” and “zoom.”

To demonstrate these animation styles, we first used UCSF Chimera [3] to generate a multi-frame PDB file of *S. cerevisiae* hexokinase 2 (*Sc*Hxk2). Specifically, we used Chimera’s “Morph Conformations” tool to capture the transition between open and closed *Sc*Hxk2 structures extracted from a recent molecular dynamics simulation [19]. We created video animations of this transition from image sequences of 48 Prot2Prot-rendered trajectory frames (Video S1).

These animations convincingly capture the *Sc*Hxk2 open-to-close transition, but the protein surfaces appear to “flicker.” This subtle artifact arises because Prot2Prot renders each frame without regard for adjacent frames. To address this issue, we used Prot2Prot to re-render the *Sc*Hxk2 trajectory to only twelve images. We then used the Real-time Intermediate Flow Estimation (RIFE) 3.1 algorithm [20], as implemented in the Flowframes software package [21], to interpolate between these twelve images. Curiously, RIFE 3.1 is also based on a neural network. The resulting animations capture the same open-to-close transition but without the flicker (Video S2). We had similar success using the commercial frame interpolation algorithm implemented in Adobe After Effects.

### Compatibility and Run Times

We have tested the Prot2Prot Web App on all major operating systems and web browsers (Table 1), including some mobile devices. The Prot2Prot model is memory intensive, and the web app will crash if run on a device with a less capable graphical processing unit (GPU). Where possible, the app detects any crash and asks the user to (1) select a smaller output-image size or (2) use the central processing unit (CPU) rather than the GPU. Rendering on the CPU is slower but also less memory restrained.

**Table 1.**
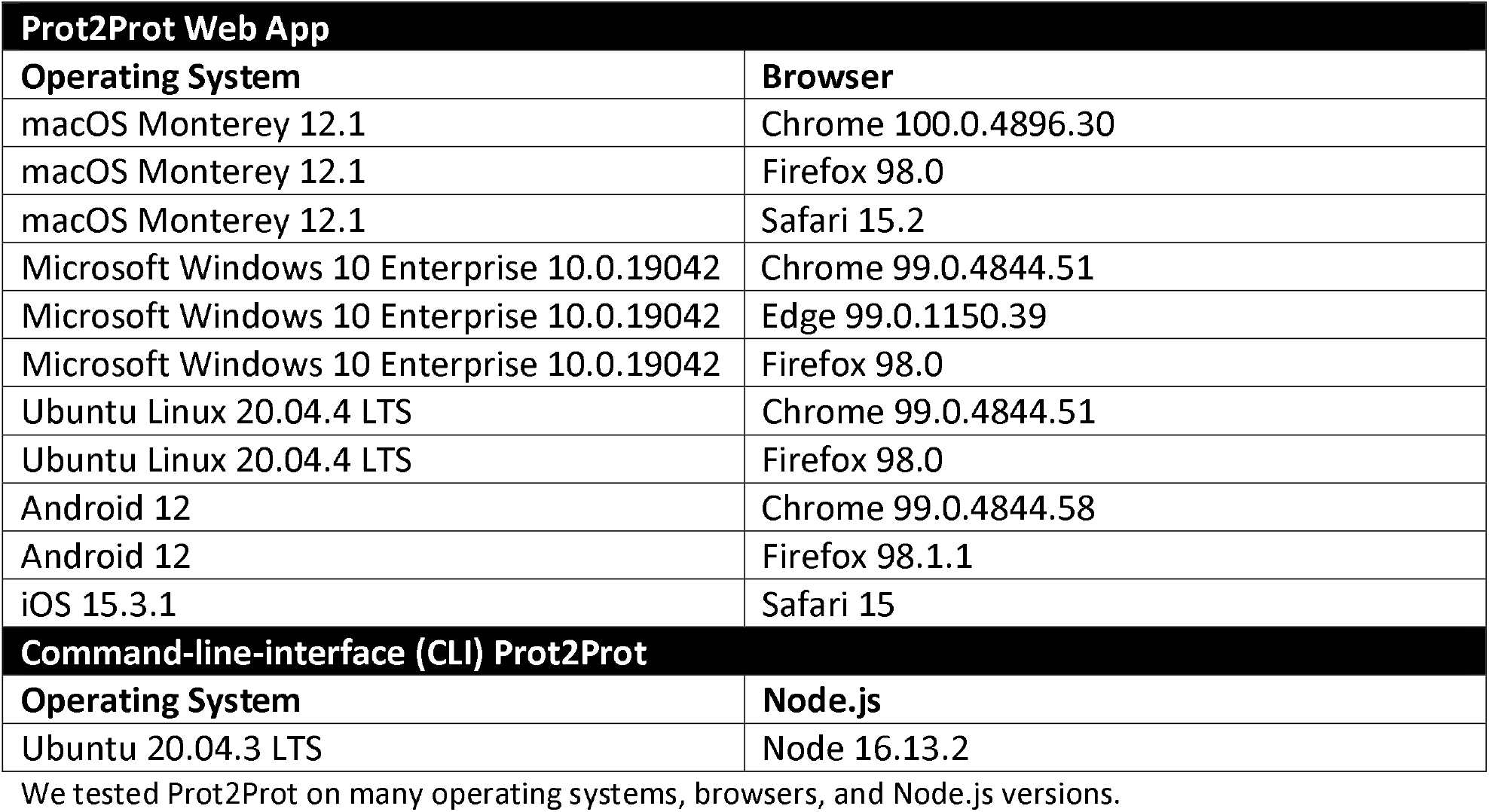
Prot2Prot compatibility.

Prot2Prot currently runs fastest on Chromium-based browsers (e.g., Google Chrome, Microsoft Edge, etc.) because these browsers support OffscreenCanvas. On other browsers (e.g., Firefox), TensorFlow.js must use the CPU to run inference rather than the GPU. Users can already enable OffscreenCanvas in Firefox via the advanced configuration preferences, suggesting future versions will enable it by default.

We tested CLI Prot2Prot on Ubuntu Linux running Node.js 16.13.2. The Node.js runtime environment is available on all major desktop operating systems, so we expect CLI Prot2Prot to be broadly compatible as well.

Aside from benefiting from broad compatibility, Prot2Prot also produces high-quality images much faster than dedicated 3D modeling programs such as Blender. To demonstrate, we rendered a test scene using Blender 3.0.0 on a MacBook Pro with an Apple M1 Max chip. The Blender Cycles path-tracing engine took roughly two minutes to generate a 1024 px x 1024 px image. In contrast, the Prot2Prot web app running on the same machine (Chrome browser) generated a similar image in only 1.2 seconds once the WebGL shaders had compiled (∼6 seconds). Rendering times vary substantially depending on the available software and hardware (e.g., GPU vs. CPU). For example, Blender 3.0.0 does not support GPU rendering on Apple hardware, and Prot2Prot does not run as quickly when using the CPU version of TensorFlow.js (as required, for example, in Firefox and Safari). But this comparison nevertheless demonstrates that Prot2Prot can dramatically accelerate photorealistic molecular visualization without requiring expertise in 3D modeling.

### Limitations

Prot2Prot is a powerful, easy-to-use tool for photorealistic protein rendering, but it has several notable limitations. First, it is generally useful only for rendering protein surfaces. We attempted to train a Prot2Prot model to generate a cartoon-like image of protein tertiary structure given a sketch of the protein backbone atoms (Figure 5, A-C). Prot2Prot often correctly identified alpha helices and beta sheets, but misclassifications were frequent. Furthermore, it depicted alpha helices as elongated blobs rather than perfect cylinders.

**Figure 5.**
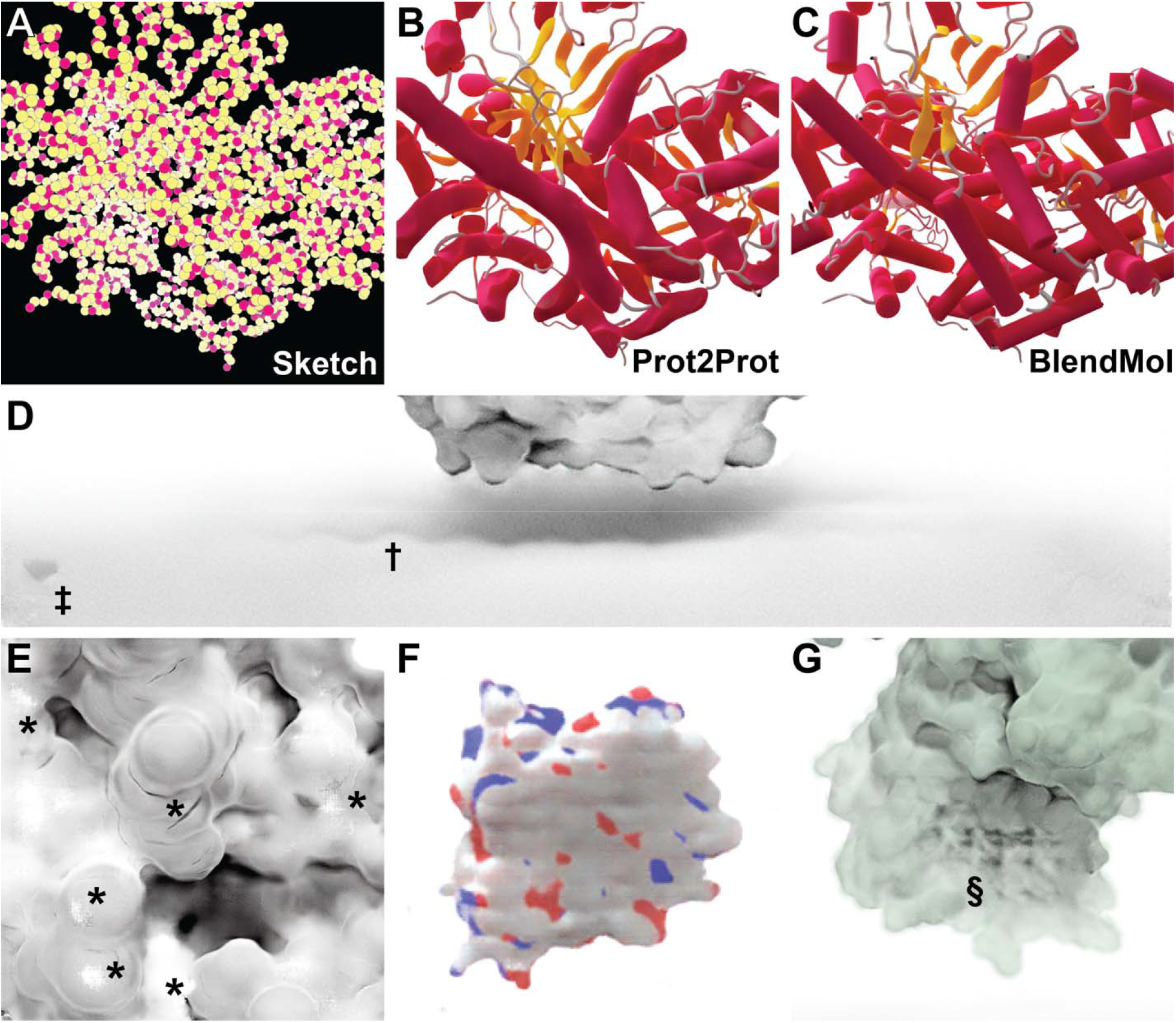
Examples of Prot2Prot shortcomings. A-B) Prot2Prot is best suited for rendering protein surfaces. It cannot accurately render a cartoon representation given a sketch of the protein backbone atoms. D) The Chalky Shadow rendering style sometimes generates shadows that are excessively wavy (†). An artifactual shadow “blob” sometimes appears in the lower-left-hand corner (‡). E) Viewing protein surfaces up close can produce artifacts (*). F) Viewing protein surfaces at great distances tends to overrepresent carbon atoms (white). G) On rare occasions protein surfaces may be subtly checkered even at intermediate distances (§). appear when rendering proteins even at intermediate distances (Figure 5G, marked with §). Rotating or scaling the molecule slightly generally eliminates these patterns.

The shadows rendered when using the Chalky Shadow style are generally impressive. Still, occasionally they appear to be more wavy than appropriate given the actual contours of the protein’s profile (Figure 5D, marked with †). Prot2Prot also sometimes renders a shadow “blob” in the lower-left-hand corner of its Chalky-Shadow output images (Figure 5D, marked with ‡). Fortunately, image cropping can easily remove this small artifact.

Prot2Prot also often struggles to correctly render protein surfaces with positioning that differs substantially from that depicted in the training images. Artifacts typically occur when proteins are very close to the virtual camera (Figure 5E, marked with *) or very distant (Figure 5F). In the case of distant proteins, Prot2Prot appears to overemphasize the contribution of carbon atoms (Figure 5F, colored in white). Finally, subtle checkered patterns occasionally

## Conclusion

The literature describes several other applications of image-to-image translation in medicine and biology. Examples include enhancing medical [22] and histopathological [23] images to facilitate diagnosis. Others have applied similar approaches to images obtained from electron [24] and fluorescence [25–27] microscopy with the goal of detecting gold nanoparticles or subcellular components. But to the best of our knowledge, these approaches have never been applied to macromolecular visualization with the goal of producing photorealistic images for scientific analysis, publication, outreach, and education.

Though the present work focuses on molecular visualization, it also suggests how machine learning algorithms can rapidly and effectively enhance scientific visualization generally. Blender specifically has been used to visualize many scientific phenomena, ranging from quantum wave functions [28] to tsunami hydrodynamics [29] to astrophysical data [30,31]. A similar approach—generating simple representations of scientific data and converting those representations to higher-quality images—could be fruitfully applied in these other domains as well.

## Acknowledgments

We thank the University of Pittsburgh’s Center for Research Computing for computer resources and Harrison Green for help with the browser implementation. This work was supported by the National Institute of General Medical Sciences of the National Institutes of Health [R01GM132353 to J.D.D.]. The content is solely the responsibility of the authors and does not necessarily represent the official views of the National Institutes of Health.

## Supporting Information

**Video S1. Prot2Prot-generated animations**. Animations depict the transition of *Sc*Hxk2 between open and closed states (3,671 atoms). A multi-frame PDB file was first generated by “morphing” between two conformations extracted from a molecular dynamics simulation. This file was then used to render 48 Prot2Prot images, which were looped when composing the video. The video also illustrates Prot2Prot’s still, rock, turntable, and zoom animations. All proteins were rendered in the Simple Surface style.

**Video S2. Prot2Prot animations with frame interpolation**. These animations were generated by applying frame interpolation (RIFE 3.1) to twelve Prot2Prot-generated images of the same *Sc*Hxk2 trajectory (3,671 atoms per frame). Top row: Proteins rendered in the Simple Surface style. Bottom row: Proteins rendered in the Chalky Shadow style. Note that the shadows update dynamically as the protein moves and that the colorization post-processing filter imparts a green tint to the protein surface.

